# Dissecting the contribution of vagal subcircuits in sepsis-induced brain dysfunctions

**DOI:** 10.1101/2024.02.14.580156

**Authors:** Lena Bourhy, Carine Moigneu, Alice Dupin, Estéban Hecquet, Jarod Levy, Tarek Sharshar, Pierre-Marie Lledo, Gabriel Lepousez

## Abstract

Sepsis, a life-threatening syndrome caused by a dysregulated host response to infection, induces a range of acute effects on the brain, including sickness behaviour and sepsis-associated encephalopathy. In addition, sepsis can lead to durable changes in neuronal circuits, resulting in long-term impairments such as post-traumatic stress disorder (PTSD). These brain dysfunctions are not directly caused by brain infection but result from peripheral inflammatory signals relayed to the brain via neural and humoral pathways. The vagal complex in the brainstem, composed of the nucleus of the solitary tract (NTS) and the area postrema, plays a crucial role in sensing and relaying these signals. Notably, the activation of the vagal complex triggers neurovegetative, neuroendocrine, and behavioural responses to infection. Chronic electrical vagus nerve stimulation has been used clinically to treat various brain disorders and is being investigated for its potential to alleviate inflammation and immune diseases through the anti-inflammatory reflex. However, a deeper understanding of the involvement of the vagus nerve and downstream brain circuits in sepsis-induced brain activation and dysfunction is needed to optimize therapeutic strategies.

To investigate the role of the vagal complex in sepsis-induced brain dysfunction, various techniques were employed to manipulate vagus nerve activity and downstream circuits in a rodent model of sepsis by caecal ligation and puncture. Subdiaphragmatic vagotomy and pharmacogenetic manipulation of NTS and nodose (i.e. vagus sensory neurons) were implemented, revealing that vagotomy effectively reduced acute brain activation, inflammatory responses, and sickness behaviour triggered by sepsis. Additionally, transient activation of NTS neurons had a significant impact on inflammatory responses, sickness behaviour, and long-term PTSD-like consequences. This study underscores the complex interplay among the vagus nerve, brain circuits, and systemic inflammation during sepsis, emphasizing the critical importance of understanding these interactions in the development of targeted therapeutic interventions.

## Introduction

Sepsis is a life-threatening syndrome characterised by a dysregulated host response to infection, referred to as cytokine storm. In addition, sepsis induces a spectrum of acute effects on the brain, ranging from sickness behaviour to encephalopathy. Sickness behaviour is considered as an adaptive response to systemic inflammatory signals triggered by infection. It is characterised by behavioural and cognitive changes (reduced locomotion and food intake, social withdrawal, attention deficits), metabolic responses (e.g. fever, glycemia), along with activation of the neuroendocrine (i.e. vasopressin and HPA axes) and neurovegetative systems (1). In addition to sickness behaviour, brain dysfunction - referred to as sepsis-associated encephalopathy (SAE) - is observed in 50 to 70% of patients. SAE is characterised by severe consciousness impairments (ranging from delirium to coma), pathological electroencephalographic changes, increased mortality, and long-term morbidity (2).

Neither sickness behaviour nor SAE are related to direct brain infection. During infection, peripheral inflammatory signals are relayed to the brain *via* neural and humoral pathways (3). In the brainstem, the vagal complex comprises different nuclei, namely the nucleus of the solitary tract (NTS) and the area postrema (AP), dedicated to sensing and relaying these neural and humoral pathways to downstream regions in the brain. The neural pathway involves local sensing of inflammatory mediators by sensory neurons of the peripheral nervous system. Among the peripheral nerves, the vagus nerve is a major relay of visceral and cardio-thoracic inflammatory messages to the central nervous system. Vagus nerve sensory neurons are pseudo-unipolar cells whose cell bodies are located within the nodose ganglia and project to the vagal complex in the brainstem. The systemic administration of the bacterial endotoxin lipopolysaccharide (LPS), cytokines, or pathogens has been shown to stimulate the vagal complex in rodents, illustrated by the increased expression of the neuronal activation marker Fos in the NTS and in the AP (4). Vagus nerve sensory fibres represent 80% of the total nerve fibres and express the nociception channel Nav1.8 by a vast majority (5,6). The remaining 20% composes the descending motor fibres of the vagus nerve. These fibres originate from cholinergic neurons in the dorsal motor nucleus (DMN) (which is also part of the vagus complex) and project directly to most organs. These fibres release neurotransmitters and neuropeptides that directly dampen immune responses at the site of infection and in immune organs such as the spleen (3,7,8). This descending response is referred to as the ‘anti-inflammatory reflex’ (4). In addition to the neural pathway, the humoral pathway includes cytokine diffusion, immune cell infiltration and endothelial cell activation in response to circulating inflammatory agents, notably through the fenestrated capillaries of brain circumventricular organs, such as the AP (9,10). In the AP, blood cytokines activate perivascular and glial cells, which relay the information to AP neurons for transmission to the NTS. The activation of the vagal complex is then transfered to different centres involved in neurovegetative (parabrachial nucleus (PBN)), neuroendocrine (paraventricular nucleus (PVN) and supra-optic nucleus (SO) of the hypothalamus) and behavioural response to infection (extended amygdala, composed of the central nucleus of the amygdala (CeA) and bed nucleus of the stria terminalis (BNST)) (11).

Chronic electrical vagus nerve stimulation (VNS) based on implantable stimulation devices is currently used in clinics to treat brain disorders such as epilepsy and depression (12,13). VNS is also tested for the alleviation of chronic inflammation and immune diseases based on the anti-inflammatory reflex (14). In rodents, experimental studies have shown that VNS effectively prevents the systemic release of pro-inflammatory cytokines, hypotension, and coagulation alterations following LPS injection (15), and may even decrease mortality (16,17). Conversely, the bilateral section of the vagus nerve abdominal branches (i.e. subdiaphragmatic vagotomy) decreases behavioural depression upon LPS injection, without affecting fever (18–20). On the other hand, the activation of a genetically defined neuronal population of the NTS can recapitulate most of the components of sickness behaviour. Thus, the mechanisms underlying the beneficial effects of VNS remain partially elucidated, due to the lack of specificity of the process, which indiscriminately modulates both sensory and motor fibres of the vagus nerve. However, the application of local ultrasound SNV at the nodose level has been found to diminish LPS-induced TNF release (21), suggesting that the modulation of sensory fibres alone may play a pivotal role in attenuating infection-induced sickness behaviour. These new medical devices, less invasive than VNS, make the procedure available to a wider range of diseases and patients (22,23). However, a better understanding of how the vagus nerve and downstream brain circuits are involved in sepsis-induced brain activation and the resulting dysfunctions is essential to decipher to what extent these techniques could be a potential therapeutic target in sepsis patients.

Interestingly, sepsis does not only impact neuronal responses during infection but can also lead to durable changes in the neuronal circuits of the brain that result in multiple long-term impairments in survivors. Recent work demonstrated that the transient activation in the first hours of sepsis of a CeA subpopulation expressing the protein kinase C delta (PKCδ+) and projecting to the ventral BNST (vBNST) was responsible for the later development of post-traumatic stress disorder (PTSD)-like behaviour in mice (24). Consequently, our hypothesis posits that if manipulation of the vagus nerve can influence brain activation during sepsis, it may likewise exert a discernible impact on long-term outcomes that remains to be characterized.

The rodent model of sepsis based on polymicrobial intra-abdominal infection after caecal ligation and puncture (CLP) mimics the clinical and pathophysiological features of human sepsis-induced cytokine storm and brain response, and leads to long-term PTSD-like behaviour in mice (24,25). To assess the role of the vagal complex in sepsis-induced brain dysfunction, we implemented different gain- and loss-of-functions techniques to manipulate vagus nerve activity and downstream circuits during CLP: subdiaphragmatic vagotomy, pharmacogenetic bidirectional manipulation of NTS and nodose neurons. Our findings revealed that vagotomy effectively reduced CLP-triggered acute activation in neurovegetative areas of the brainstem, curbed inflammatory responses, and alleviated sickness behaviour. While manipulating both NTS and Nodose neurons bidirectionally influenced Fos expression in the CeA, only NTS pharmacogenetic transient activation had a significant impact on CeA PKCδ-expressing neurons, CLP-induced inflammatory responses, sickness behaviour, and long-term PTSD-like consequences.

## Methods

### Animals

Adult (2-5 months old) wild-type male and female C57Bl/6JRj mice (Janvier labs, Le Genest-Saint-Isle, France) and male and female Nav1.8 Cre mice (*Scn10a*^*tm2(cre)Jwo*^; Jackson labs; (26)) were housed under a 12-h light/dark cycle, with dry food and water available ad libitum. All procedures were consistent with the European Union guidelines (EU Directive 2010/63/EU) for animal experiments and were reviewed and approved by the Animal Welfare Committee of the Institut Pasteur (Projects numbers: 2015-004, 2013-0086, dap180018 and dap200025).

### Subdiaphragmatic vagotomy

Mice were treated for analgesia and rehydration 30 minutes before surgery with subcutaneous (SC) injection of buprenorphine (0.1mg/Kg, Vetergesic 0.3mg/mL, Ceva Santé Animale, Libourne, France) and saline (NaCl 0.9%). The vagotomy surgery (VagX) was achieved under general anaesthesia (Isoflurane, 4% for induction in the inhalation chamber, then maintained during the surgery at 1.5% with 98.5% oxygen). Mice were placed on a heating pad to maintain body temperature at 37°C during the procedure. Absence of reaction to leg and tail pinch was checked before incision of the abdominal wall previously shaved and cleaned with betadine (soap and solution). The abdomen cavity was opened, and the stomach was isolated from the liver and the gut using surgical tensioners and humidified compresses. Under a binocular loop, the gastric anterior branch of the vagus nerve was identified at the junction between the oesophagus and the stomach, grasped with surgical forceps and cut on a segment of 2 to 3 mm with surgical scissors. The same protocol was achieved for the gastric posterior branch of the vagus nerve. Compresses and tensors were then removed, and the abdominal muscles and peritoneum were sutured in two separate plans. Buprenorphine and saline SC injections were performed every 12 hours until complete recovery. Control animals (Ctl) underwent the same anaesthetic, analgesic, and surgical protocol that VagX mice except for the grasp and the section of the vagus nerve branches.

### Cholecystokinin (CCK) test

Two weeks after vagotomy, mice were isolated and left 24 hours in their new cage for habituation. At the end of the habituation period, they were fasted for another 24 hours and then injected with CCK (40 ug/kg diluted in ammonia injected intraperitoneally (IP)) and immediately offered a new food pellet for 1 hour. A group of naïve mice following the same protocol but injected with PBS was added as a control condition of basal food intake after 24-hour fasting. The pellet weight was measured before and one hour after injection. The difference of the pellet weight was normalised to the mouse weight.

### Cecal ligation and puncture (CLP) surgery

Mice were treated for analgesia and rehydration 30 minutes before surgery with SC injection of buprenorphine (0.1mg/Kg, Ceva Santé Animale, Libourne, France) and saline (NaCl 0.9%). The CLP surgery (∼10 min) was achieved under general anaesthesia (Isoflurane, 4% for induction in the inhalation chamber, then maintained during the surgery at 1.5% with 98.5% oxygen).

Mice were placed on a heating pad to maintain body temperature at 37°C during the procedure. Absence of reaction to leg and tail pinch was checked before incision of the abdominal wall previously cleaned with ethanol 70%, and then incision of the peritoneum. Caecum was exposed and a loose ligation was achieved at its external third with Mersilk 4.0 (Ethicon, Cincinatti, USA). Two transfixing punctures were performed avoiding blood vessels with a 21G needle. Faeces were expressed and spread on caecum before suturing abdominal muscles and peritoneum in two separate plans. Buprenorphine and saline (same concentrations) SC injections were then performed every 12 hours until complete recovery. We evaluated sepsis severity every 12 hours for 72 hours using the sepsis score, which evaluates the clinical signs of local (abdominal spasm) and systemic reaction to sepsis (rectal temperature, fur erection, abnormal breathing), the behavioural changes (faeces cleaning, presence of peri-ocular dried eye drop, incomplete palpebral opening, spontaneous activity in the cage, the escape attempt after tail grabbing) and muscular weakness (ability to grip to forceps, body tone, difficulty to walk). Each item of the scale was scored 0 or 1, with a total score ranging from 0 (normal) to 12 (highly severe). For ethical reasons, mice with a sepsis score ≥6 and a body temperature ≤35.3°C were euthanized. CLP mice were compared to Sham animals that only underwent a laparotomy procedure with the same anaesthetic and analgesic protocol than CLP mice. CLP and sham surgery were performed in the morning (Zeitgeiber time 3-5).

### Stereotaxic injections

Mice were transported to the room where the surgery was performed and underwent a SC injection of buprenorphine (49,5ug/kg, Vetergesic 0.3mg/mL, Ceva Santé Animale, Libourne, France) and saline (NaCl 0.9%) 30 minutes before anaesthesia. Mice were anesthetized with an intra-peritoneal (IP) injection of xylazine (5mg/Kg, Rompun 2%, Bayer, Leverkusen, Germany), and ketamine (125mg/Kg, Imalgene 1000, Mérial, Lyon, France) and then placed on a heating pad to maintain body temperature at 37°C. Stereotaxic injections of viral vectors in adult mice were performed as previously described (24). Briefly, the animal’s head was inserted into a stereotaxic frame (David Kopf Instruments). Following local anaesthesia (lidocaine, Xylovet, France), the animal’s head was shaved, the scalp sterilized with an iodine solution and cut to reveal the skull. Small craniotomies were drilled and the tip of the pulled glass pipettes (tip diameter, ∼30-50 μm) of the injection system (Nanoinject II, Drummond) were slowly lowered to the target coordinates. Some AAV preparations were mixed before injection. In mix 1, AAV1-hSyn-Cre-WPRE (University of Pennsylvania, 1.10e13 vg/mL, 1:20 of final mix) was mixed with AAV5-hSyn-DIO-mCherry-WPRE (Addgene 50459, 2.5.10^13^ vg/mL, diluted 1:5 in Saline 0.9%, 19:20 of final mix). In mix 2, AAV1-hSyn-Cre-WPRE (University of Pennsylvania, 1.10e13vg/mL, 1:20 of final mix) was mixed with AAV5-hSyn-DIO-hM3D(Gq)-mCherry (Addgene 44361, 6×10^12^ vg/mL, not diluted, 19:20 of final mix). Wild-type animals were then injected whether with mix 1 or mix 2 or AAV9-CamKII-GCaMP6f-WPRE-SV40 (Addgene, 100834, 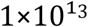 vg/mL) or AAV9-CamKII-hM4D(Gi)-mCherry (construct provided by Bryan Roth, production by the Vector core of the Gene Therapy Laboratory of Nantes, INSERM UMR1089, 1.1x10^13^ vg/mL) in the NTS (from bregma, AP: -7.4, ML: ±0.35, DV from bregma: -5.0, 50nL injected bilaterally, speed 2nL/s). pAAV5-hSyn-DIO-mCherry (Addgene, 50459, 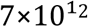 vg/mL), pAAV5-hSyn-DIO-hM4D(Gi)-mCherry (Addgene, 44362, 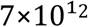 vg/mL) or pAAV5-hSyn-DIO-hM3D(Gq)-mCherry (Addgene, 44361, 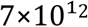 vg/mL) were injected in Nav1.8 Cre animals in in the NTS (from bregma, AP: -7.4, ML: ±0.35, DV from bregma: -5.1 and -5,4, 50nL injected bilaterally at each depth, speed 2nL/s). At the end of the injection, the skin was sutured and then sterilized with iodine solution. The animal was injected SC with meloxicam (2,5mg/Kg, Metacam 5mg/mL, Boehringer Ingelheimwas, Ingelheim-Am-Rhein, Germany) and then transferred back to its home cage. Animals in which post-hoc histological examination showed that viral injection were not in the correct location were excluded from analysis.

### Pharmacology

Clozapine-*N*-oxide (CNO) (C0832; Sigma-Aldrich: Stock solution in DMSO at 100mg/mLresuspended in 0.9% NaCl saline, 1 mg/kg) was injected intraperitoneally at time 0 hours (H0), H6, H12 and H24 according to the experiment.

### Contextual fear conditioning

Behavioural assays started 16 days after CLP. Mice were transported in their home cage to the experiment room 45 min prior experiments. A 100-lux light was used, and mice were tracked with a JVC lowlux camera (Yokohama, Japan) fixed to the ceiling above the arenas. Mice were individually placed in a homemade electrifiable grid floor Plexiglas arena (20x15cm) contained in a sound-proof box connected to a constant-current shock generator (Supertech instruments, Switzerland) and equipped with a camera and an audio speaker. Before each conditioning session, the cage was washed with Surfa’safe® premium (Anios, Lille). After a 3-minute habituation period, mice underwent two sequences of a 28-second 2.5 kHz tone (conditional stimulus (CS), repetition frequency: 440 Hz, 75 Db) followed by a 2-s electric shock (0.4 mA, unconditional stimulus (US)) separated by a 15-s silence period, then followed by a 1-min resting period. Movement following shock was monitored through a camera placed above the arena. Fear recall was performed 24 h after conditioning. The conditioning and the contextual recall environment were identical. Mice behaviour was recorded for 3 minutes, and live freezing time was evaluated. Freezing was defined as the absence of movement except for breathing.

### Immunohistochemistry

Under deep anaesthesia (xylazine 20 mg/kg, ketamine 100 mg/kg), both nodose ganglia were exposed by making an incision along the ventral surface of the neck and blunt dissection, then extracted, and fixed with 4% Paraformaldehyde overnight, before being stored in phosphate buffered saline (PBS) with 30% sucrose. Mice were next euthanized by an intra-cardiac infusion of 0.9% NaCl then 4% Paraformaldehyde. Brains were extracted with forceps after incision of the skin and medial section of the skull with fine scissors. Brains were post-fixed with 4% Paraformaldehyde for 2 hours, then rinsed with PBS before being stored in PBS with 30% sucrose. Cryostat sections of nodose ganglia (16μm) were first dehydrated with 100% ethanol, then rinsed in 0.1 m PBS and then processed for single- or double-labeling immunofluorescence. Brains were coronally cut into 60 μm thick sections with a freezing microtome (Leica). The sections of interest were stored in PBS with 0.01% Azide. The immuno-labelling protocol begun with a permeabilization step using PBS with 0.2% Triton X-100 (PBST 0.2%) and 10% Normal Donkey Serum (NDS, Abcam, Ab7475) blocking agent for 2 hours under constant agitation. Primary antibodies diluted according to the supplier’s recommendations were then incubated with PBST 0.2%, NDS 10% and azide 0.01%, at 4°C with constant agitation for 2 days. Primary antibody used was Fos (1:1000, Rabbit, Abcam, Ab190289) and mCherry (1:4000, Goat, Sicgen, AB0081). After three 5-minute washes with PBS, sections were incubated with either Alexa Fluor 488, 568 or 647–conjugated secondary antibodies (Invitrogen, ThermoFisher) with PBST 0.2%, NDS 2% and DAPI (0.01%) at room temperature with constant agitation for 2 hours. After three 5-minute washes with PBS, slices were mounted on slides with a mounting medium (FluoroMount-G, Interchim). Image acquisition was carried out with a digital slide scanner (Axio Scan, ZEISS). Fos quantification was achieved using the “spot detection” function of ICY software (Institut Pasteur, France Bioimaging), allowing to count the number of Fos+ cells in brain areas previously drawn on the slides’ images via the ROI (Region of interest) function of the same software. The brain areas were identified according to the reference *Allen brain* atlas.

### Cytokine and chemokine multiplex

Blood was collected directly from the heart under terminal anaesthesia right before euthanizing in 1.5 mL tubes (Microvet tubes), serum was obtained with centrifugation for 5 min at 10000rpm, immediately frozen and stored at -20°C until assay. Cytokines and chemokines were measured using Bio-Plex Pro Mouse Cytokine 23-Plex Immunoassay (M60009RDPD, BioRad, USA) following manufacturer’s recommendations, which allow measurement of Eotaxin, G-CSF, GM-CSF, IFN-γ, IL-1α, IL-1β, IL-2, IL-3, IL-4, IL-5, IL-6, IL-9, IL-10, IL-12 (p40), IL-12 (p70), IL-13, IL-17A, KC, MCP-1 (MCAF), MIP-1α, MIP-1β, RANTES, TNF. Cytokines’ concentrations were normalised to the total protein concentration in each serum aliquot. Cytokines which did not reach the level of detection were not included in the analysis.

### Statistical analysis

All experiments and data analyses were achieved blindly, unless otherwise stated. In the figure legends are indicated the number of subjects used in each experimental condition and the data are expressed as mean ±SD. Two-sided statistical analyses were performed with GraphPad Prism 6.0. No statistical methods were used to pre-determine sample size, or to randomise. Samples normality was previously tested using D’Agostino & Pearson omnibus normality test. For analysis of variance tests, we used ANOVA tests with multiple comparisons. Outliers were identified using Grubbs’ Method (α=0.05) and then removed. Correlations were assessed by calculating Pearson r coefficients. Statistical significance was set at *P < 0.05, **P < 0.01, ***P < 0.001, ****P < 0.0001.

### Data availability

The datasets generated during the current study are available from the corresponding authors upon reasonable request.

## Results

### Multi-scale models to manipulate the activity of distinct vagal system circuits during sepsis

Research studies conducted on both rodents and humans have emphasised that sepsis triggers a transient and significant neuronal activation in various brain regions. To mimic sepsis in mice, we used the validated murine model of sepsis induced by peritonitis after CLP. We quantified the neuronal activation marker Fos in the brain and showed that Fos expression was significantly increased at 6 hours post-CLP (H6) in the main output region of the vagus nerve, i.e. the NTS (Fig.1e-f). We therefore hypothesise that the vagus nerve could play a pivotal role in relaying sepsis signals to the brain. To assess this hypothesis, we implemented three different techniques aiming at manipulating the activity of distinct vagal system subcircuits during CLP (27).

**Figure 1.**
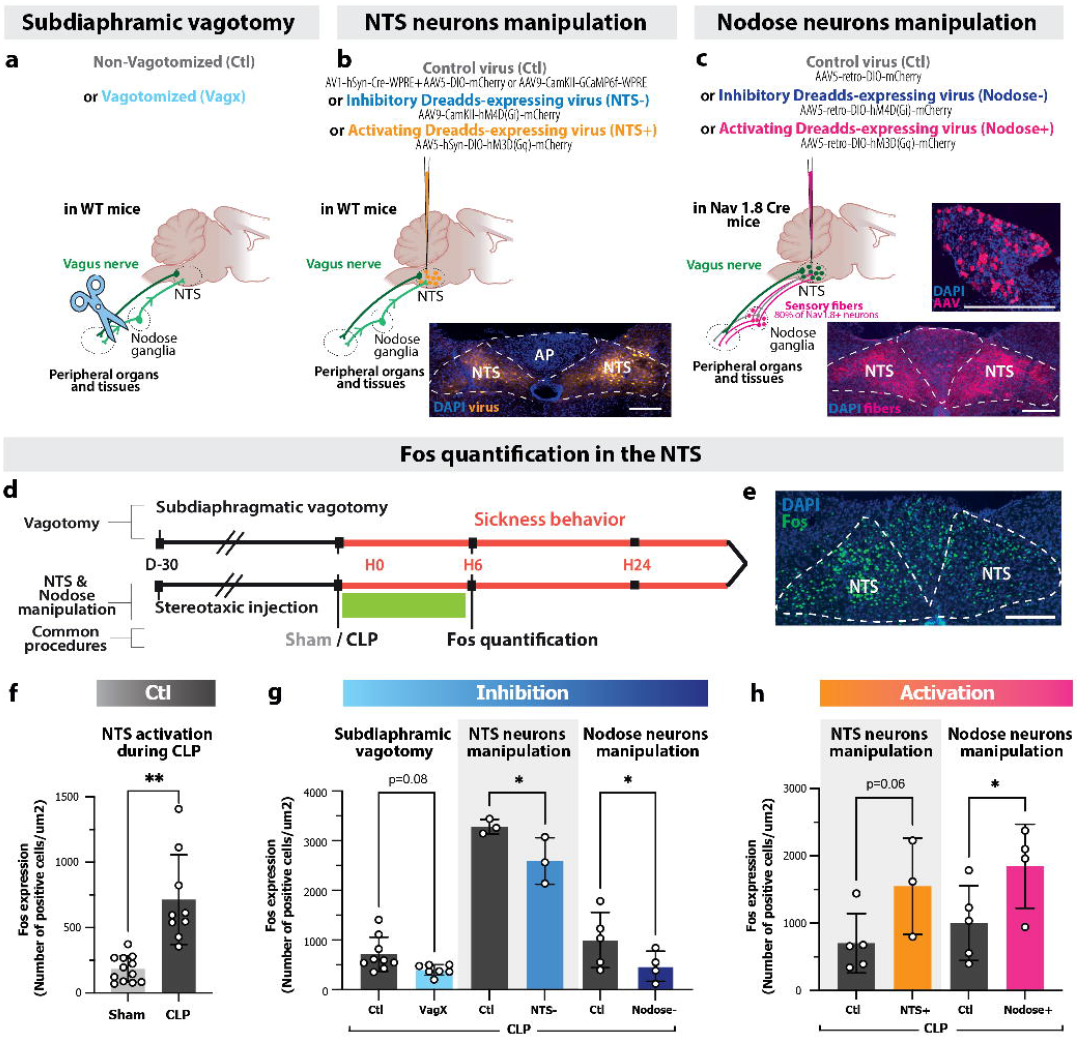
Validation of vagus complex manipulation models. **a**. In the subdiaphragmatic vagotomy model, both branches of the vagus nerve were surgically cut at the abdominal level (Vagx), 30 days before CLP surgery (D-30). **b**. In the NTS neurons manipulation model, a viral vector expressing either a control protein (Ctl) or an inhibitory (NTS-) / activating (NTS+) DREADD was injected directly in the NTS at D-30. CNO was then injected during the first 24 hours following CLP to allow the transient inhibition/activation of NTS neurons. **c**. In the Nodose neurons manipulation model, a retrograde viral vector expressing either a Cre-dependent control protein (Ctl) or an inhibitory (Nodose-) / activating (Nodose+) DREADD was injected in the NTS of Nav-1.8 Cre mice at D-30, allowing with CNO the specific and transient inhibition/activation of the sensory fibres of the vagus nerve (i.e. Nodose ganglia neurons) during the first 24 hours following CLP. **d**. Timeline of the experiment according to the model. **e**. Fos expression in the NTS. **f**. In control condition (without any manipulation of the vagus nerve neurons), CLP induces a strong neuronal activation in the NTS compared to Sham animals, highlighted by an increase Fos expression at 6 hours (H6) (n_Ctl-Sham_=12, n_Ctl-CLP_=9, Unpaired t-test **P=0.0015). **g**. Vagus nerve inhibition exhibited a dampening effect on the CLP-induced NTS activation at H6 in the three model we tested (Two-way ANOVA, effect of inhibition ***P=0.0009, multiple comparisons: VAG: n_Ctl-CLP_ =9, n_(-)-CLP_=7, P=0.083; NTS: n=3, *P=0,023; Nodose: n_Ctl-CLP_ =5, n_(-)-CLP_=4, *P=0.032) **h**. NTS or Nodose activation amplified Fos expression at H6 post-CLP in the NTS (Two-way ANOVA, effect of activation *P=0.010, multiple comparisons: NTS: n_Ctl_ =5, n_(+)_=3, P=0.063; Nodose: n_Ctl_ =5, n_(+)_=4, *P=0.046). NTS: Nucleus of the tractus solitarii, Scale=100um.

In the first model, mice underwent subdiaphragmatic sections of the abdominal branches of the vagus nerve to suppress vagal communication at the abdominal level, while leaving the thoracic vagal innervation intact (Fig.1a). Vagotomized mice (Vagx) were compared to non-vagotomized animals (Ctl) that underwent the same anaesthetic and surgical procedure except for the vagus nerve sections. To validate this model, two weeks after surgery, mice were fasted for 24 hours and then injected with the satiating agent cholecystokinin (CCK), whose receptors are mainly located on the abdominal vagus nerve fibres (SFig.1). The results indicated that, within the first hour following CCK injection, Ctl mice—unlike Vagx mice—significantly reduced their food intake compared to naive mice injected with PBS, thereby confirming the disruption of the CCK vagal relays in Vagx animals. 30 days post-vagotomy, mice had fully recovered from the surgery and underwent CLP (Fig.1d). Vagotomy demonstrated efficiency, as evidenced by the reduced Fos expression induced by CLP in the NTS at H6 in Vagx animals compared to Ctl (Fig.1g).

To expand on this first loss-of-function approach and improve the time-specificity of vagus nerve manipulation, we used pharmacogenetic tools. In a first model, we targeted the vagus nerve output neurons by injecting directly in the NTS a control virus (Ctl) or a Designer Receptor Exclusively Activated by Designer Drugs (DREADD) virus, expressing either the inhibitory receptor hM4D(Gi) (NTS-) or the activating receptor hM3D(Gq) (NTS+) in wild type animals (Fig.1b). To narrow down neuronal specificity, we also targeted in another model the expression of DREADD viruses specifically in the vagus nerve sensory neurons using genetically modified Nav1.8 Cre mice, which express Cre-recombinase enzyme exclusively in Nav1.8 neurons, i.e. 80% of the sensory neurons of the vagus nerve (28) (Fig.1c). These mice were also injected in the NTS but this time with a retrograde Cre-dependant virus expressing either a control virus (Ctl) or the inhibitory (Nodose-) or activating (Nodose+) DREADD receptor. Conditional recombination enabled the precise manipulation exclusively of the sensory neurons of the vagus nerve projecting to the NTS, whose cell bodies are located in the nodose ganglia (Fig.1c). DREADDs are triggered exclusively in presence of Clozapine N-oxide (CNO); 30 days after stereotaxic injections, CNO was administered to mice within the first 24 hours post-CLP, allowing transient and reversible inhibition/activation of the targeted neurons solely during this specific time frame (Fig.1d). These pharmacogenetic manipulations demonstrated efficiency, as evidenced by the respective reduction or increase in Fos expression in the NTS at H6 post-CLP (Fig.1g-h).

These findings also underscored the considerable variability in Fos expression levels, influenced by factors such as the experimental model, mouse strain, or the AAV promoter employed (Ca^2+^/calmodulin-dependent protein kinase II (CamKII) for NTS- and human synapsin (hSyn) in NTS+ and Nodose-/+) (Fig.1g-h). To enhance clarity and facilitate comprehension, the forthcoming data will be presented as normalised to each model Ctl levels.

Together, these results highlighted 3 different approaches to efficiently manipulate vagal complex inputs and outputs with increasing temporal and neuronal specificity. Consequently, we employed these approaches to assess the impact of such silencing/enhancing of vagus nerve relays on CLP-induced activation in other brain regions.

### Vagus complex manipulation impacts CLP-driven transient activation of the brainstem and the amygdala

To decipher the role of the vagus nerve in sepsis-induced brain activation pattern, we evaluated the effect of the modulations induced by the above approaches over H6 CLP-induced Fos expression in brain regions downstream to the vagal complex. As previously shown, CLP increased neuronal activity in brain areas involved in the neurovegetative (PBN), neuroendocrine (PVN, SO) and emotional (CeA, vBNST) responses to infection (Fig.2a, SFig.2). Notably, CLP-induced Fos expression in the NTS was strongly correlated with Fos expression in these other brain regions, as evidenced by Pearson r calculations (Fig.2g, SFig.2).

**Figure 2.**
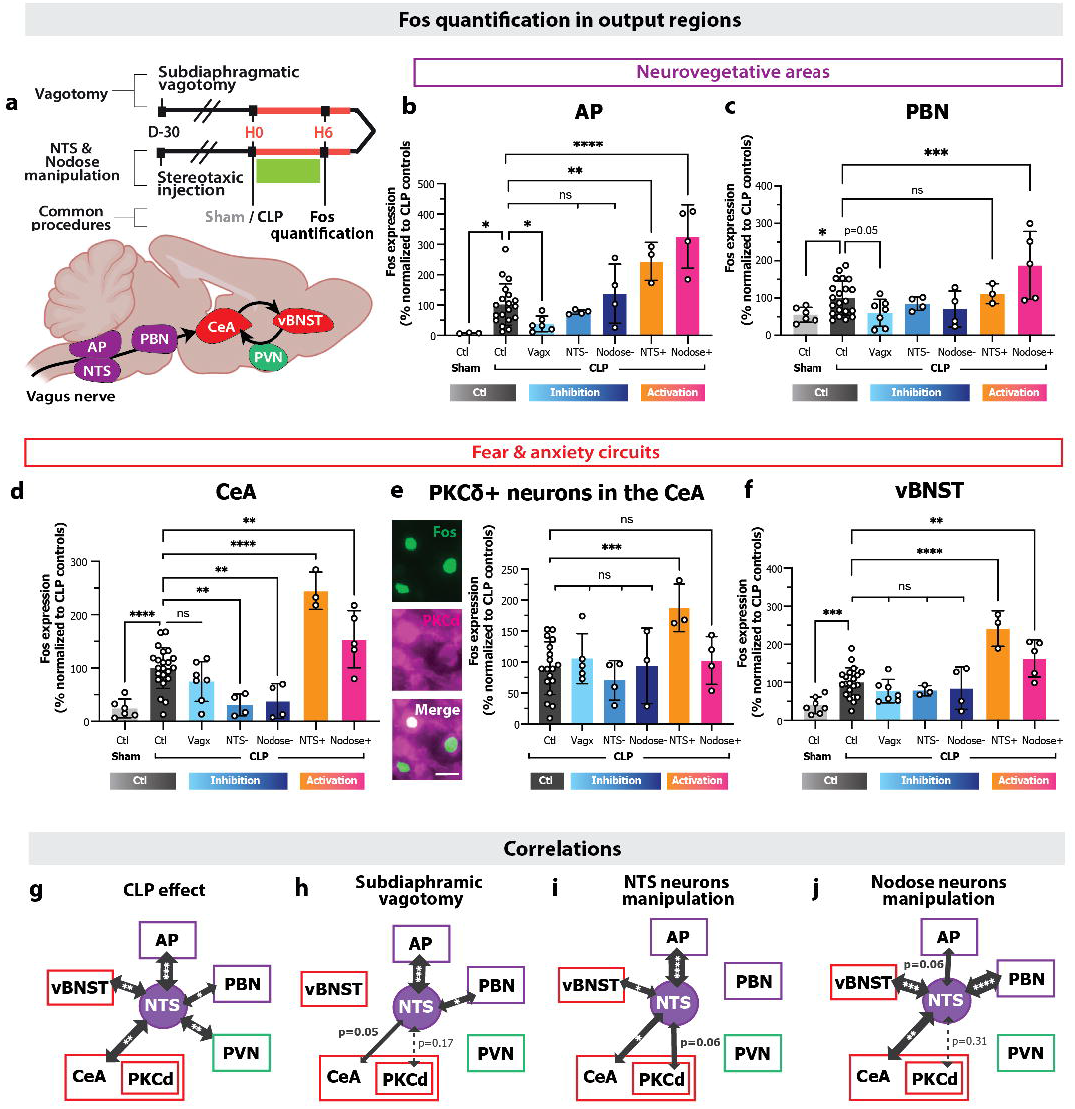
Impact of vagus complex manipulation on CLP-induced transient brain activation. **a**. Timeline of the experiment according to the models, and schematic of the regions of interest screened in the Fos quantification process. **b**.**c**. Vagus nerve inhibition dampened the CLP-induced activation in the AP and the PBN at H6 only in the vagotomy model, whereas NTS or Nodose activation increased Fos expression (One-way ANOVA, AP: n_Ctl-Sham_ =3, n_Ctl-CLP_ =19, n_VagX-CLP_=6, n_NTS(-)-CLP_=4, n_Nodose(-)-CLP_=4, n_NTS(+)-CLP_=3, n_Nodose(+)-CLP_=4; multiple comparisons: Ctl-Sham vs Ctl-CLP: *P=0.027, Ctl-CLP vs Vagx-CLP:*P=0.047, Ctl-CLP vs NTS(-)-CLP: P=0.500, Ctl-CLP vs Nodose(-)-CLP: P=0.330, Ctl-CLP vs NTS(+)-CLP: **P=0.001, Ctl-CLP vs Nodose(+)-CLP: ****P<0.0001, PBN: n_Ctl-Sham_ =6, n_Ctl-CLP_ =21, n_VagX-CLP_=7, n_NTS(-)-CLP_=4, n_Nodose(-)-CLP_=4, n_NTS(+)-CLP_=3, n_Nodose(+)-CLP_=5; multiple comparisons: Ctl-Sham vs Ctl-CLP: *P=0.043, Ctl-CLP vs Vagx-CLP: P=0.058, Ctl-CLP vs NTS(-)-CLP: P=0.550, Ctl-CLP vs Nodose(-)-CLP: P=0.252, Ctl-CLP vs NTS(+)-CLP: P=0.682, Ctl-CLP vs Nodose(+)-CLP: ***P=0.0005). **d**.**e**. NTS and Nodose neurons manipulation modulated bidirectionally Fos expression in the CeA. However, only the activation of the NTS neurons had an impact on the CLP-induced activation of the CeA PKC^™^+ subpopulation (CeA: One-way ANOVA, n_Ctl-Sham_ =6, n_Ctl-CLP_ =22, n_VagX-CLP_=7, n_NTS(-)- CLP_=4, n_Nodose(-)-CLP_=4, n_NTS(+)-CLP_=3, n_Nodose(+)-CLP_=5; multiple comparisons: Ctl-Sham vs Ctl-CLP: ****P<0.0001, Ctl-CLP vs Vagx-CLP: P=0.118, Ctl-CLP vs NTS(-)-CLP: **P=0.001, Ctl-CLP vs Nodose(-)-CLP: **P=0.003, Ctl-CLP vs NTS(+)-CLP: ****P<0.0001, Ctl-CLP vs Nodose(+)-CLP: **P=0.005; PKC^™^+ neurons in the CeA: One-way ANOVA, n_Ctl-CLP_ =18, n_VagX-CLP_=5, n_NTS(-)-CLP_=4, n_Nodose(-)-CLP_=3, n_NTS(+)-CLP_=3, n_Nodose(+)-CLP_=4; multiple comparisons: Ctl-CLP vs Vagx-CLP: P=0.475, Ctl-CLP vs NTS(-)-CLP: P=0.405, Ctl-CLP vs Nodose(-)-CLP: P=0.889, Ctl-CLP vs NTS(+)-CLP: ***P=0.0008, Ctl-CLP vs Nodose(+)-CLP: P=0.597). **f**. Inhibition of vagus nerve had no effect on the CLP-induced vBNST activation whereas the transient pharmacogenetic activation amplified Fos expression (One-way ANOVA, n_Ctl-Sham_ =7, n_Ctl-CLP_ =22, n_VagX-CLP_=7, n_NTS(-)-CLP_=3, n_Nodose(-)-CLP_=4, n_NTS(+)-CLP_=3, n_Nodose(+)-CLP_=5; multiple comparisons: Ctl-Sham vs Ctl-CLP: ***P=0.0007, Ctl-CLP vs Vagx-CLP: P=0.168, Ctl-CLP vs NTS(-)-CLP: P=0.358, Ctl-CLP vs Nodose(-)-CLP: P=0.466, Ctl-CLP vs NTS(+)-CLP: ****P<0.0001, Ctl-CLP vs Nodose(+)-CLP: **P=0.002). **g-h**. Correlation between Fos expression in the NTS and in other brain regions according to the type of manipulation implemented (Pearson correlations, n and p values in SFigure 2.f). Scale=20⌈m. AP: area postrema, PBN: parabrachial nucleus, CeA: central nucleus of the amygdala, PKC^™^: protein kinase C ^™^, vBNST: ventral bed nucleus of the stria terminalis.

Vagotomy emerged as the sole inhibition approach resulting in a reduction in CLP-induced activation in the neurovegetative areas located in the brainstem, such as the AP and the PBN (major output of the vagal complex) (Fig.2b-c). However, vagotomy did not present any effect on the downstream regions of the brainstem such as the hypothalamus (PVN) and the extended amygdala (CeA, BNST) (Fig.2d,f, SFig2). On the other hand, DREADD-driven NTS or Nodose neurons transient inhibition during CLP was not sufficient to significantly decrease CLP-induced circuit of activation, except for the CeA (Fig.2b-f).

Conversely, the activation of NTS or Nodose neurons during CLP globally amplified Fos expression at H6 post-CLP in all the regions aforementioned. Intriguingly, despite the bidirectional modulation of Fos expression in the CeA induced by NTS and Nodose neurons manipulation, only the activation of NTS neurons demonstrated an impact on the CeA PKCδ-expressing neurons, a subpopulation that has been shown to be responsible for the post-sepsis PTSD-like behaviour (24) (Fig.2e).

To elucidate the distinct consequences of vagus complex manipulations, we depicted correlations in Fos expression between the NTS and various brain regions activated during CLP, as per the implemented model. We employed Pearson correlation coefficient calculations for analysis (Fig. 2h-j, SFig.2f). Notably, across all models, Fos expression in the NTS showed no significant correlation with Fos expression in the neuroendocrine regions. Vagotomy, however, revealed a robust correlation exclusively between Fos expression in the NTS and that in the neurovegetative areas within the brainstem. Beyond the brainstem, manipulations of both NTS and Nodose neurons demonstrated a robust correlation between NTS Fos expression and extended amygdala Fos expression, with an increasing strength from the NTS to the Nodose model. Interestingly, when manipulating NTS neurons alone, a noteworthy correlation emerged between NTS Fos expression and the specific PKCδ-positive subpopulation of the CeA (Fig. 2i).

This finding highlights the intricate interplay and specificity of Fos expression patterns in response to different vagus complex manipulations during CLP. Notably, these results indicate that vagus complex manipulation can influence sepsis-induced activation of the brainstem and the extended amygdala. Given the pivotal role of these regions in the immune and behavioural responses to sepsis, we hypothesised that such manipulations could also exert effects on the sickness behaviour exhibited by CLP mice during the acute phase of the disease.

### Vagal complex manipulation can modulate CLP-induced sickness behaviour

To evaluate sickness behaviour after CLP, a sepsis score taking into consideration clinical and behavioural criteria was set to regularly evaluate over time mice symptoms of infection. We confirmed that CLP was associated with a significantly increased sepsis score between H12 and H24 following surgery compared to Sham animals (Fig.3a-b). The vagotomy and the DREADD-driven activation of the NTS neurons were the only manipulations that influenced the sepsis score, mitigating or exacerbating it, respectively (Fig. 3b).

**Figure 3.**
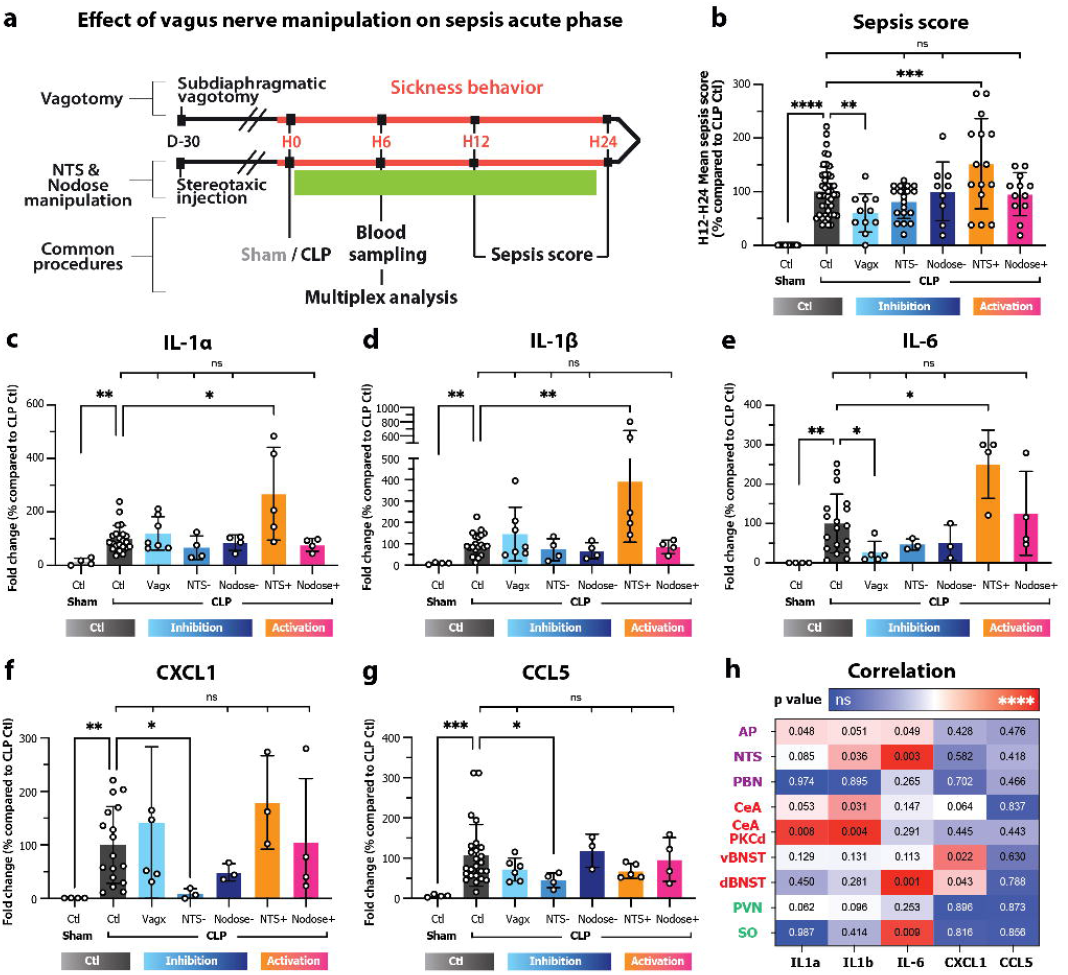
Effects of vagus complex manipulation on CLP-induced systemic inflammatory response. **a**. Timeline of the experiment. **b**. Sepsis score, that recapitulates the CLP-induced sickness behaviour, was assessed between H12 and H24. Subdiaphragmatic vagotomy improved sickness behaviour in CLP animals whereas NTS neurons activation significantly worsened it. The other manipulations had no effect (One-way ANOVA, n_Ctl-Sham_ =28, n_Ctl-CLP_ =44, n_VagX-CLP_=11, n_NTS(-)-CLP_=19, n_Nodose(-)-CLP_=9, n_NTS(+)-CLP_=15, n_Nodose(+)-CLP_=12; multiple comparisons: Ctl-Sham vs Ctl-CLP: *****P<0.0001, Ctl-CLP vs Vagx-CLP: **P=0.010, Ctl-CLP vs NTS(-)-CLP: P=0.117, Ctl-CLP vs Nodose(-)-CLP: P=0.975, Ctl-CLP vs NTS(+)-CLP: ***P=0.0002, Ctl-CLP vs Nodose(+)-CLP: P=0.752). **c-e**. Vagus nerve manipulation did not impact the CLP-induced secretion of the pro-inflammatory cytokines IL-1⟨, IL-1® and IL-6 except for NTS neurons transient activation which significantly increased their blood concentrations. Subdiaphragmatic vagotomy did present a dampening effect, only with IL-6 (IL-1⟨, Kruskal-Wallis test, n_Ctl-Sham_ =4, n_Ctl-CLP_ =22, n_VagX-CLP_=7, n_NTS(-)-CLP_=4, n_Nodose(-)-CLP_=4, n_NTS(+)-CLP_=5, n_Nodose(+)-CLP_=4; multiple comparisons: Ctl-Sham vs Ctl-CLP: **P=0.002, Ctl-CLP vs Vagx-CLP: P=0.719, Ctl-CLP vs NTS(-)-CLP: P=0.257, Ctl-CLP vs Nodose(-)-CLP: P=0.852, Ctl-CLP vs NTS(+)-CLP: *P=0.028, Ctl-CLP vs Nodose(+)-CLP: P=0.572; IL-1®, Kruskal-Wallis test, n=n_IL-1⟨_ except, n_Ctl-CLP_ =20; multiple comparisons: Ctl-Sham vs Ctl-CLP: **P=0.003, Ctl-CLP vs Vagx-CLP: P=0.651, Ctl-CLP vs NTS(-)-CLP: P=0.434, Ctl-CLP vs Nodose(-)-CLP: P=0.282, Ctl-CLP vs NTS(+)-CLP: **P=0.028, Ctl-CLP vs Nodose(+)-CLP: P=0.572; IL-6, Kruskal-Wallis test, n_Ctl-Sham_ =4, n_Ctl-CLP_ =18, n_VagX-CLP_=5, n_NTS(-)-CLP_=3, n_Nodose(-)-CLP_=3, n_NTS(+)-CLP_=4, n_Nodose(+)-CLP_=4; multiple comparisons: Ctl-Sham vs Ctl-CLP: **P=0.001, Ctl-CLP vs Vagx-CLP:*P=0.042, Ctl-CLP vs NTS(-)-CLP: P=0.405, Ctl-CLP vs Nodose(-)-CLP: P=0.380, Ctl-CLP vs NTS(+)-CLP: *P=0.036, Ctl-CLP vs Nodose(+)-CLP: P=0.666). **f-g**. Vagus nerve manipulation did not impact the CLP-induced secretion of the chemokines CXCL1 and CCL5 except transient NTS inhibition that significantly decreased their blood concentrations (CXCL1, Kruskal-Wallis test, n_Ctl-Sham_ =4, n_Ctl-CLP_ =17, n_VagX-CLP_=6, n_NTS(-)-CLP_=3, n_Nodose(-)-CLP_=3, n_NTS(+)-CLP_=3, n_Nodose(+)-CLP_=4; multiple comparisons: Ctl-Sham vs Ctl-CLP: **P=0.002, Ctl-CLP vs Vagx-CLP: P=0.699, Ctl-CLP vs NTS(-)-CLP: *P=0.014, Ctl-CLP vs Nodose(-)-CLP: P=0.465, Ctl-CLP vs NTS(+)-CLP: P=0.238, Ctl-CLP vs Nodose(+)-CLP:P=0.900; CCL5, Kruskal-Wallis test, n_Ctl-Sham_ =4, n_Ctl-CLP_ =25, n_VagX-CLP_=6, n_NTS(-)-CLP_=4, n_Nodose(-)-CLP_=3, n_NTS(+)-CLP_=5, n_Nodose(+)-CLP_=4; multiple comparisons: Ctl-Sham vs Ctl-CLP:***P=0.0005, Ctl-CLP vs Vagx-CLP: P=0.363, Ctl-CLP vs NTS(-)-CLP: *P=0.034, Ctl-CLP vs Nodose(-)-CLP: P=0.348, Ctl-CLP vs NTS(+)-CLP: P=0.345, Ctl-CLP vs Nodose(+)-CLP:P=0.927). **h**. Correlation between cytokine & chemokine concentrations and Fos expression in the brain at H6 post-CLP. (Pearson correlation, n=14, illustrated with corresponding p-values, Pearson r coefficients in SFigure 3.o). PVN: paraventricular nucleus of the hypothalamus, dBNST: dorsal BNST, SO: supraoptic nucleus.

To complement these findings, we collected blood samples from the tested animals at H6 and conducted a comprehensive cytokine profile analysis using Multiplex (Fig.3a). We confirmed that CLP induced a severe systemic peripheral inflammatory response characterised by the increased concentrations of circulating cytokines and chemokines (Fig.3c-g, SFig.3). The transient activation of NTS neurons accentuated the CLP-induced release of the pro-inflammatory cytokines IL-1α, IL-1β and IL-6 (Fig.3c-e). On the other hand, VAG significantly dampened IL-6 response to CLP, aligning with the observed sepsis score in these subjects (Fig.3b-e.). Additionally, NTS neurons transient inhibition decreased the CLP-induced secretion of the proinflammatory chemokines CX-CL1 and CCL-5 (Fig.3f-g).

We then illustrated the correlation matrix between cytokine systemic concentrations and Fos expression in the brain at H6 (Fig.3h, SFig.3o). Circulating IL-6 levels exhibited a robust correlation with Fos expression in the NTS. Conversely, concentrations of IL-1α and IL-1β were more closely correlated with Fos expression in the CeA, with a notable specificity observed in the CeA PKCδ-positive subpopulation.

Considering that CeA PKCδ-positive subpopulation activation during CLP has been shown to be responsible for the post-sepsis PTSD-like behaviour (24), we questioned whether transient vagus nerve manipulation during the acute phase of the disease could influence the long-term PTSD observed in sepsis survivors.

### Transient NTS neurons activation during CLP worsens long-term PTSD-like behaviour

To assess PTSD-like behaviour in these animals, we evaluated fear memory using the contextual fear conditioning paradigm (FC) at day 16 (D16) post-CLP, a time point when animals exhibited no lingering signs of sickness (Fig. 4a). The fear conditioning test involved pairing an electric foot shock with a specific context (29,30). Twenty-four hours after the acquisition phase, we evaluated fear memory retrieval by monitoring context-induced freezing behaviour.

**Figure 4.**
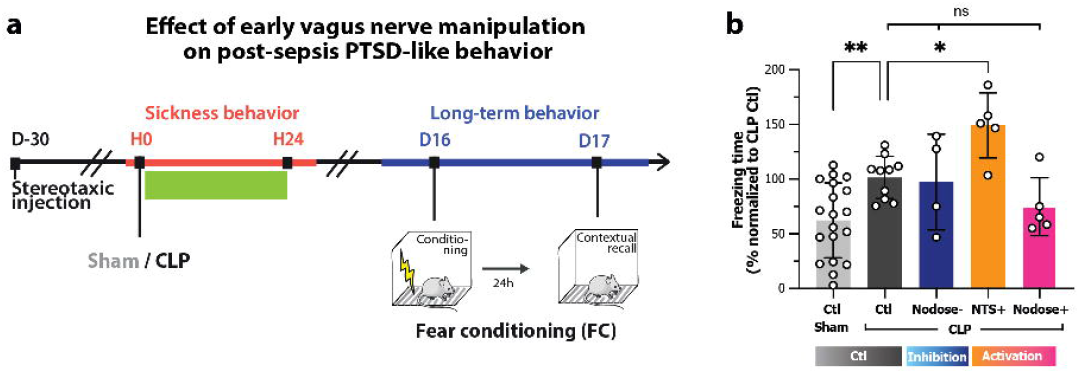
NTS activation during CLP worsened long-term PTSD-like phenotype. **a**. Timeline of the experiment. **b**. Increased freezing behaviour in CLP animals compared to Sham was amplified when NTS neurons were activated during the first 24 hours following CLP, whereas other manipulations had no effect (One-way ANOVA, n_Ctl-Sham_ =19, n_Ctl-CLP_ =10, n_Nodose(-)-CLP_=4, n_NTS(+)-CLP_=5, n_Nodose(+)-CLP_=5; multiple comparisons: Ctl-Sham vs Ctl-CLP: **P=0.010, Ctl-CLP vs Nodose(-)-CLP: P=0.815, Ctl-CLP vs NTS(+)-CLP: *P=0.024, Ctl-CLP vs Nodose(+)-CLP: P=0.232).

While transient vagal complex manipulations during CLP showed no long-term impact on contextual FC behaviour in Sham animals, they unveiled notable differences in freezing behaviour among Ctl animals at D17 post-CLP (SFig.4). Notably, Ctl-CLP animals in NTS+ and Nodose-/+ models exhibited a significant increase of freezing time during FC recall compared to Ctl-Sham (SFig.4b). However, counterparts that underwent vagotomy and NTS-manipulation did not exhibit a similar increase in freezing time. The variations observed in freezing behaviour among control animals may be attributed to factors like the specific experimental model, mouse strain, or the AAV promoter employed in the study.

Consequently, we opted to refine our assessment, focusing only on models that already displayed a significantly increased freezing behaviour in Ctl-CLP animals compared to Ctl-Sham animals at D17. Notably, DREADD-driven activation of the NTS solely within the initial 24 hours following CLP was sufficient to exacerbate post-sepsis PTSD-like fear expression in the FC test at D17 (Fig. 4b). These findings imply that temporary manipulation of the vagus nerve may influence not only sepsis-induced short-term but also long-term behaviour.

## Discussion

The major findings of this study include the following: 1) We developed efficient gain- and loss-of-function techniques to manipulate vagus nerve and downstream vagal complex activity during CLP. 2) Vagotomy was the sole loss-of-function model that effectively reduced CLP-triggered activation in neurovegetative areas of the brainstem, dampened inflammatory responses, and alleviated acute sickness behaviour. 3) While pharmacogenetic manipulation of both NTS and Nodose neurons bidirectionally influenced Fos expression in the CeA, only NTS neurons transient activation had a significant impact on the CLP-induced activation of CeA PKCδ+ neurons CeA and the later PTSD-like consequences.

Our observations revealed that vagotomy improved the acute response to CLP, in contrast to the transient inhibition of the NTS or nodose neurons (Fig.2-3). In addition, our results showed that only vagotomy was significantly decreasing CLP-induced activation in the AP (Fig.2b). This discrepancy can be attributed to the intricate network of the vagus nerve, which not only projects to the NTS but also to the AP. For instance, vagal sensory neurons that express neuropeptide Y receptor Y2 and originate from the heart ventricular wall have been shown to specifically connect to the AP, but not the NTS (31). Interestingly, the AP is constituted of different neuronal subpopulations. For instance, a subpopulation expressing Adcyap1 in the NTS and AP is necessary and sufficient to induce sickness symptoms, specifically anorexia and lethargy (but nor fever) in mice injected with LPS (20). These different subpopulation are also distinguishable by their projections which connect, independently of the NTS, to various regions activated during CLP: AP excitatory neurons project to many brain regions, including PBN, PVN, and nearby NTS regions, whereas inhibitory projections are largely confined within the AP itself, and to a lesser extent within proximal NTS regions, but are not observed in the PBN (32) (Fig.2c). Significantly, only vagotomy exhibited a decrease in the CLP-induced activation of the PBN. This implies that inhibiting NTS alone is insufficient to mitigate neuronal activation in the PBN, owing to the vagal pathway through the AP, vagal collaterals outside of the abdominal cavity and probably the activation of AP excitatory neurons during CLP via the humoral pathways. Indeed, the AP plays a pivotal role not only in the neural pathway but also in the humoral pathway (1). Its fenestrated capillaries allow the direct passage of inflammatory mediators into the brain parenchyma, potentially triggering robust AP activation independently of vagus nerve stimulation. Recognizing CLP as a severe inflammatory model, it is likely that pathways beyond the vagus nerve, specifically the humoral pathway, may contribute to a consistent response during CLP (33). To achieve a thorough comprehension of its contributions to the intricate network of responses, the next crucial steps involve manipulating both AP neurons and nodose neurons projecting to the AP.

In contrast to brainstem regions, manipulation of NTS or nodose neurons exhibited a capacity to modulate Fos expression in the CeA, while vagotomy did not induce a similar effect (Fig.2d). Notably, only the activation of NTS neurons led to an increase in Fos expression in the CeA PKCδ+ neurons, a subpopulation implicated in the later development of PTSD during CLP (Fig.2e). Accordingly, the transient activation of NTS neurons emerged as the exclusive model capable of amplifying PTSD-like behaviour on D17 in CLP mice (Fig.4). These findings underscore the notion that the diverse models employed can selectively target distinct populations of vagal complex neurons, each projecting to specific brain circuits. Numerous studies have emphasised the heterogeneity of NTS neuron subtypes, their projections, and functions. For example, NTS epinephrine neurons co-expressing neuropeptide Y have been identified as stimulators of feeding, while activation of NTS norepinephrine neurons has been shown to suppress feeding (34). He et al. demonstrated that the development of depression-like behaviours under chronic pain is specifically linked to the activation of direct glutamatergic projections from the NTS to somatostatin-expressing neurons in the CeA (35). Additionally, a recent study revealed that proenkephalin-expressing cells within the CeA receive direct inputs from the NTS (36). Hence, it is plausible to hypothesise that various types of NTS neurons may be activated during sepsis, projecting either directly or indirectly to distinct subpopulations within the CeA.

Interestingly, the inflammatory signature associated with the activation of these different circuits appears to differ. Notably, CLP-induced NTS activation is correlated to IL-6 release whereas PKCd+ CeA activation is correlated to IL-1α and β release (Fig.3h). It has been demonstrated that systemic pro-inflammatory injections predominantly activate specific neuronal circuits in the brain (37,38). Intriguingly, the activation of specific brain neurons has also been found to elicit distinct peripheral immune responses (39) and to recruit anti-inflammatory pathways via the release of corticosterone (38) or the activation of the cholinergic anti-inflammatory pathway (15). This suggests the possibility that each presumed circuit may exert specific pro-inflammatory effects or is selectively activated in the presence of particular cytokines. However, the current experimental design does not allow us to discern whether this correlation is explained by a causal effect from the brain to the periphery or vice versa.

In our study, we were unable to clearly establish the initial distinction in long-term fear responses between Ctl-Sham and Ctl-CLP mice in the vagotomy and NTS-models (9) (SFig.4). Non-vagotomized or NTS-Ctl-CLP mice exhibited a freezing response to fear conditioning comparable to that of Ctl-CLP mice from the other models. However, non-vagotomized and NTS-Ctl-Sham mice displayed an unusually heightened freezing behaviour. This PTSD-like response observed even in non-vagotomized sham animals may be attributed to the initial laparotomy undergone by both vagotomized and non-vagotomized animals (to expose the abdominal branches of the vagus nerve). This procedure is repeated 4 weeks later during the Sham/CLP surgeries. Despite stringent aseptic precautions for both surgeries, we cannot rule out the possibility that inflammation resulting from abdominal exploration and potential local contamination might induce an immune response akin to an initial mild sepsis. This could explain the heightened freezing behaviour observed in Sham animals. Moreover, due to technical considerations, the NTS-model was the only one employing an AAV with the CamKII promoter, in contrast to the hSyn promoter used in NTS+ and Nodose-/+ models. The expression of this virus may exhibit higher toxicity compared to the others, potentially contributing to the heightened PTSD-like behaviour observed in Sham animals.

Finally, our results raise questions about the therapeutic interest of VNS in sepsis patients. Specifically, our study revealed that chemogenetic stimulation of the NTS resulted in heightened sickness behaviour, inflammatory response, and subsequent development of a PTSD-like syndrome. These outcomes cast doubt on the potential beneficial effects of such manipulations and warrant further consideration in the context of therapeutic interventions for sepsis. However, the intricate nature of vagus nerve electrical activation patterns has been demonstrated in previous studies (38–41). To pinpoint the precise activation patterns associated with different conditions in sepsis mice, *in vivo* recordings of vagus nerve activity would be essential. This technique holds promise for distinguishing between ‘pathological’ and ‘beneficial’ electrical patterns, paving the way for the development of specific VNS protocols with therapeutic implications. Intriguingly, our earlier research indicated that administering the neuro-modulatory drug levetiracetam (LEV) intraperitoneally during the initial 48 hours post-CLP selectively reduced neuronal activation in the vagal complex and extended amygdala (22). Notably, only intraperitoneal, not intracerebroventricular, administration of LEV was effective in mitigating sepsis-induced PTSD-like behaviour, suggesting a preferential action on the peripheral nervous system. These findings collectively propose that LEV may alleviate post-sepsis PTSD by influencing the vagus nerve. Conducting *in vivo* recordings of vagus nerve activity during CLP after LEV administration could be a valuable first step in identifying potential ‘beneficial’ electrical patterns within the vagus nerve.

