## Supplementary figures for "Dissecting the contribution of vagal subcircuits in sepsis-induced brain dysfunctions"

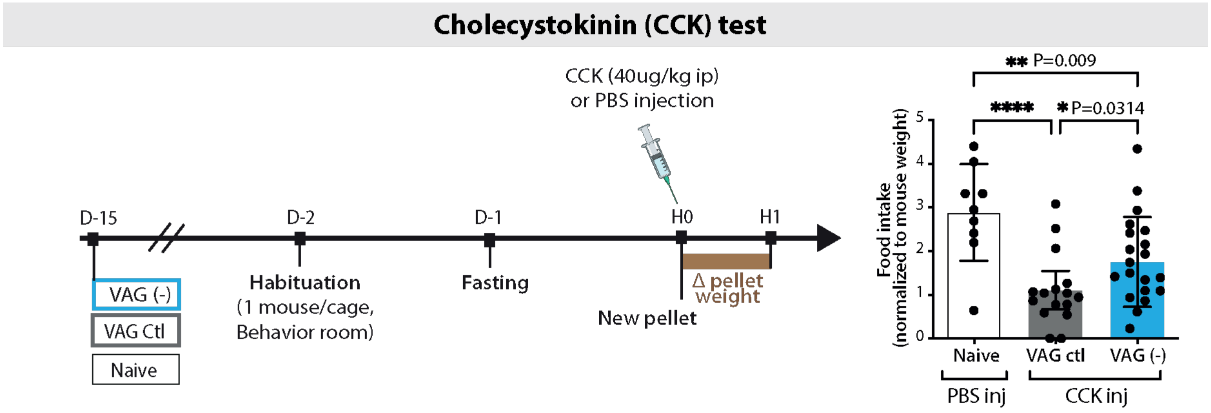


SFig1

**Supplementary figure 1.** Vagotomy was validated using the CCK test**.** After 24-hour fasting, food intake after CCK injection was significantly less decreased in vagotomized animals compared to Sham (One-way ANOVA: $n_{naive}$=9, $n_{VAGctl}$=16, $n_{VAGx}$=20, ***P=0.0003).

### Fos quantification in the brain

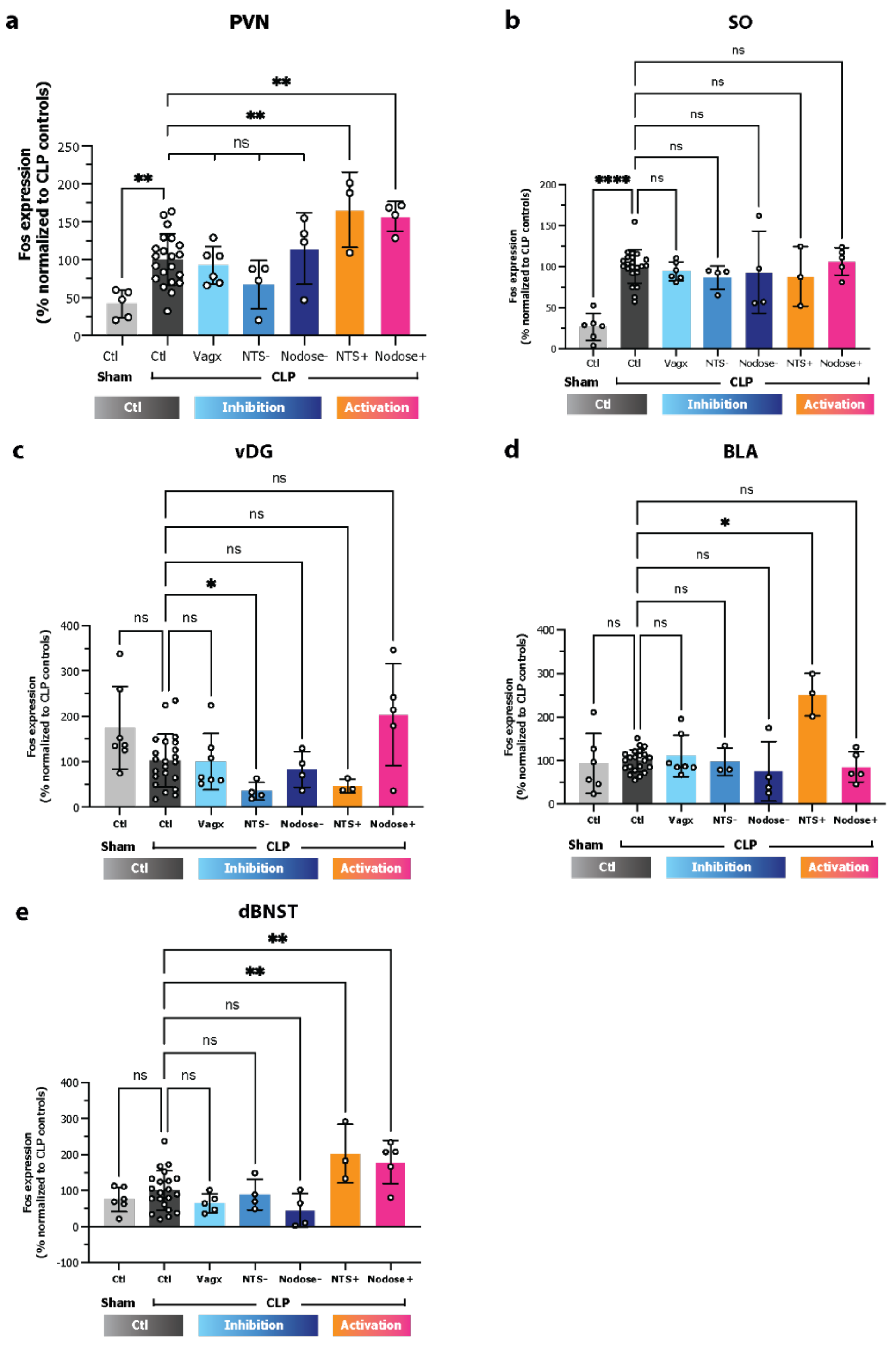


**f** **Correlations**

**CLP effect**

| | NTSvs.AP | NTSvs.PBN | NTSvs.CeA | NTSvs.vBNST | NTSvs.dBNST | NTSvs.PVN | NTSvs.SO | NTSvs.vDG | NTSvs.BLA |
|---|---|---|---|---|---|---|---|---|---|
| number of pairs | 9 | 11 | 12 | 13 | 11 | 11 | 12 | 11 | 12 |
| P value | 0.002 | 0.012 | 0.0012 | 0.0038 | 0.4159 | 0.002 | 0.0102 | 0.587 | 0.2029 |

**Subdiaphragmatic vagotomy**

| | NTSvs.AP | NTSvs.PBN | NTSvs.CeA | NTSvs.vBNST | NTSvs.dBNST | NTSvs.PVN | NTSvs.SO | NTSvs.vDG | NTSvs.BLA | NTSvs.PKCd Cea |
|---|---|---|---|---|---|---|---|---|---|---|
| number of pairs | 13 | 14 | 15 | 15 | 12 | 14 | 14 | 15 | 13 | 5 |
| P value | 0.0003 | 0.0293 | 0.059 | 0.5648 | 0.8004 | 0.8486 | 0.1984 | 0.8274 | 0.4321 | 0.171 |

**NTS neurons manipulation**

| | NTSvs.AP | NTSvs.PBN | NTSvs.CeA | NTSvs.vBNST | NTSvs.dBNST | NTSvs.PVN | NTSvs.SO | NTSvs.vDG | NTSvs.BLA | NTSvs.CeA PKCd |
|---|---|---|---|---|---|---|---|---|---|---|
| number of pairs | 14 | 15 | 15 | 15 | 15 | 15 | 14 | 14 | 12 | 15 |
| P value | <0.0001 | 0.1781 | 0.0426 | 0.0457 | 0.0415 | 0.5318 | 0.9021 | 0.8168 | 0.0441 | 0.0639 |

**Nodose neurons manipulation**

| | NTSvs.AP | NTSvs.PBN | NTSvs.CeA | NTSvs.vBNST | NTSvs.dBNST | NTSvs.PVN | NTSvs.SO | NTSvs.vDG | NTSvs.BLA | NTSvs.CeA PKCd |
|---|---|---|---|---|---|---|---|---|---|---|
| number of pairs | 12 | 13 | 13 | 13 | 13 | 11 | 13 | 13 | 13 | 7 |
| P value | 0.0693 | <0.0001 | 0.0018 | 0.0003 | 0.001 | 0.2544 | 0.3163 | 0.5566 | 0.9634 | 0.3127 |

**Supplementary figure 2. a-e.** Brain Fos expression at H6 post-CLP in our three different models (One-way ANOVA and multiple comparison test). **f.** Correlation of Fos expression in the NTS and other brain regions according to the model of vagus nerve manipulation (Pearson correlations, number of pairs and P values). PVN: paraventricular nucleus of the hypothalamus, SO: Supraoptic nucleus, vDG: ventral dendrite gyrus of the hippocampus, BLA: basolateral nucleus of the amygdala, dBNST: dorsal bed nucleus of the stria terminalis.

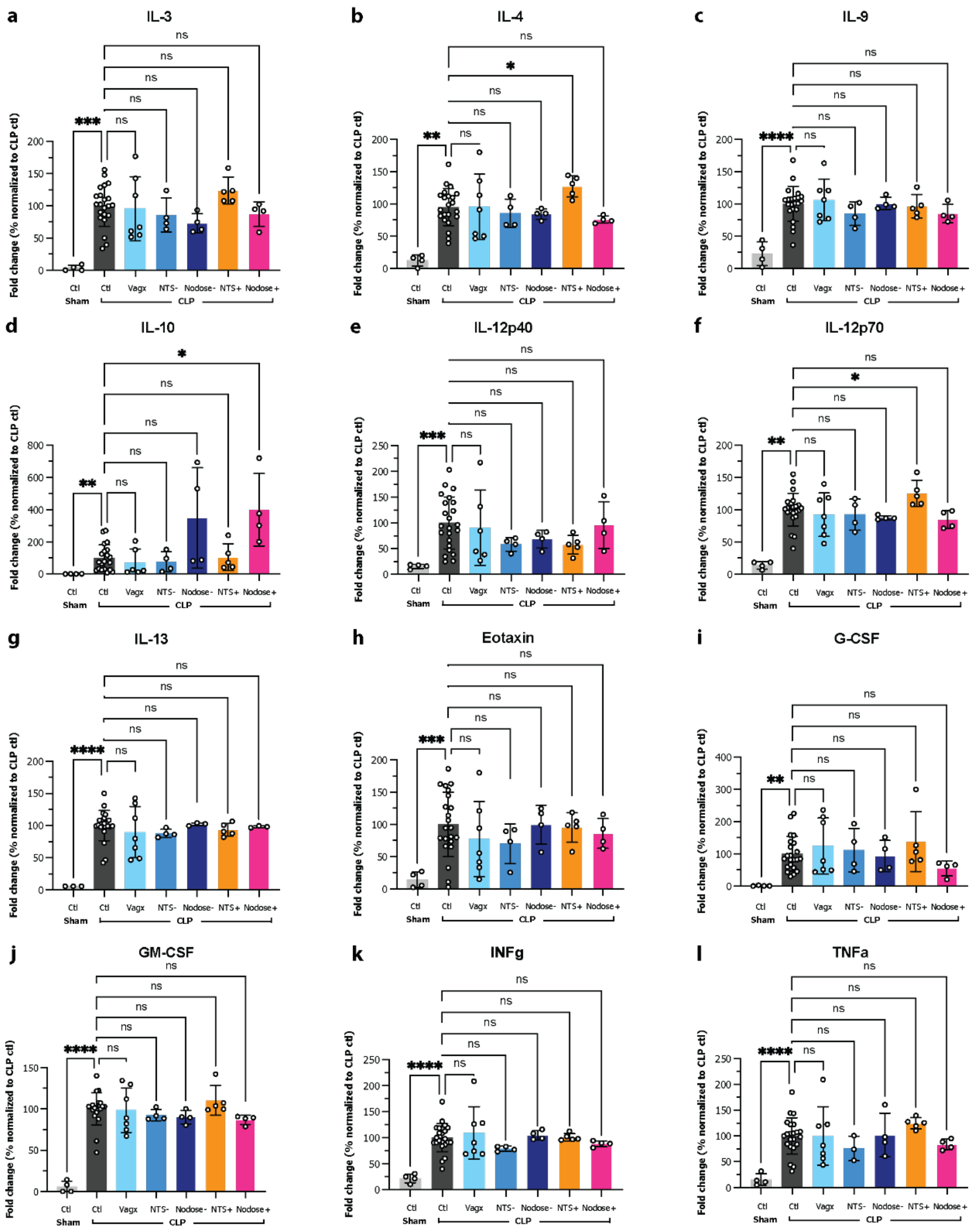


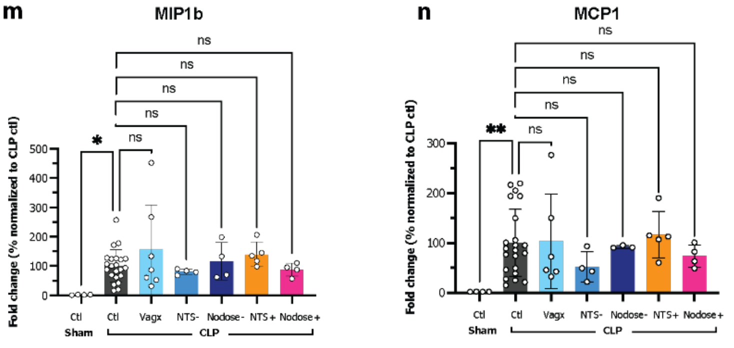


**o** Correlation - Pearson r coefficients

| | IL1a | IL1b | IL-6 | CXCL1 | CCL5 |
|---|---|---|---|---|---|
| AP | 0.518 | 0.513 | 0.535 | 0.283 | 0.200 |
| NTS | 0.459 | 0.543 | 0.724 | 0.199 | 0.226 |
| PBN | -0.009 | -0.037 | 0.320 | 0.139 | 0.204 |
| BLA | 0.241 | 0.154 | 0.155 | 0.545 | -0.118 |
| CeA | 0.508 | 0.558 | 0.409 | 0.604 | -0.058 |
| vBNST | 0.410 | 0.408 | 0.443 | 0.706 | -0.136 |
| dBNST | 0.211 | 0.298 | 0.773 | 0.647 | -0.076 |
| vDG | -0.072 | -0.009 | -0.148 | 0.487 | 0.008 |
| PVN | 0.493 | 0.445 | 0.327 | 0.048 | 0.045 |
| SO | -0.005 | -0.228 | -0.670 | 0.085 | 0.051 |
| CeA PKCd | 0.659 | 0.697 | 0.304 | 0.273 | -0.214 |

**SFig3**

**Supplementary figure 3. a-n.** Cytokine and chemokine blood concentrations at H6 post-CLP in our three different models (One-way ANOVA and multiple comparison test). **o.** Correlation of Fos expression in the brain and cytokine/chemokine concentrations at H6 post-CLP (Pearson r coefficients).

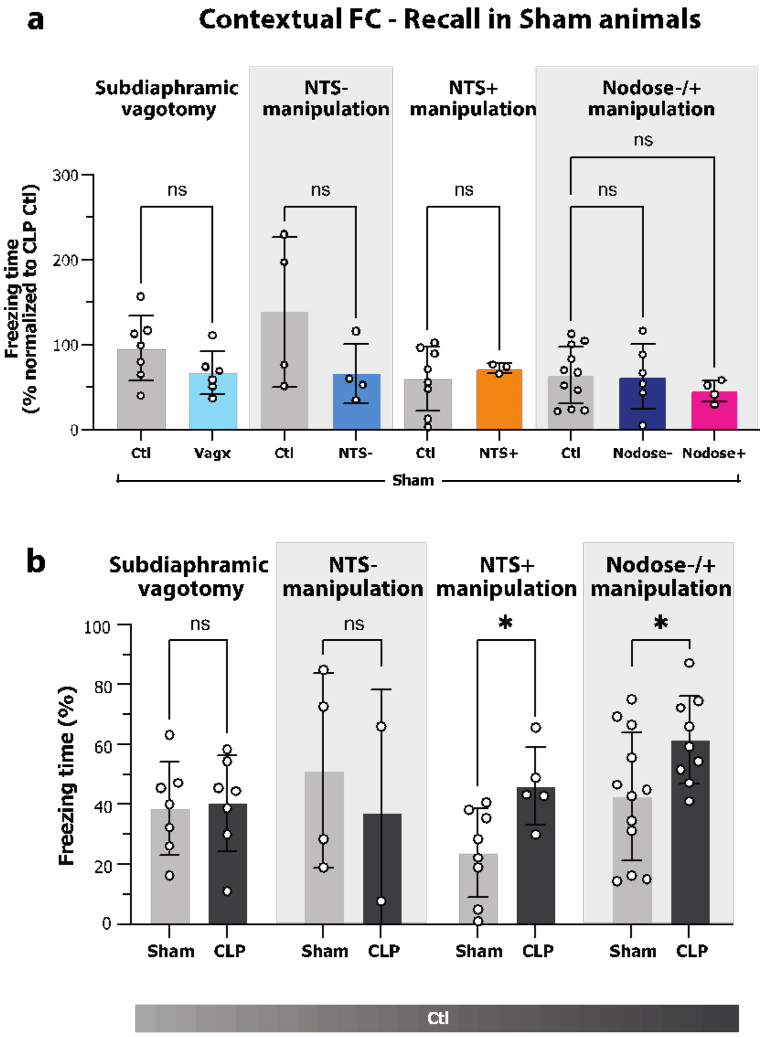


### SFig4

**Supplementary figure 4. a.** Contextual fear conditioning response at D17 in Sham animals according to the model of vagus nerve manipulation (Two-way ANOVA and multiple comparison test). **b.** Response to fear conditioning at D17 post-CLP in control animals was different according to the model implemented. CLP animals presented an increased freezing time compared to Sham animals in NTS+ and Nodose +/- controls whereas no PTSD-like behaviour was observed in the vagotomy and NTS- controls (Two-way ANOVA, group effect P=0.097, multiple comparisons: VAG: n=7, P=0.860; NTS-manipulation: $n_{Ctl\text{-}Sham}$=3, $n_{Ctl\text{-}CLP}$ =2, P=0.857; NTS+ manipulation: $n_{Ctl\text{-}Sham}$=8, $n_{Ctl\text{-}CLP}$ =5, *P=0.039; Nodose-/+ manipulation: $n_{Ctl\text{-}Sham}$=12, $n_{Ctl\text{-}CLP}$ =9, *P=0.025).
